# Exploiting O-GlcNAc transferase promiscuity to dissect site-specific O-GlcNAcylation

**DOI:** 10.1101/2023.07.31.551258

**Authors:** Conor W. Mitchell, Sergio Galan Bartual, Andrew T. Ferenbach, Carsten Scavenius, Daan M. F. van Aalten

**Author notes:** These authors contributed equally to this work (Conor W. Mitchell, Sergio Galan Bartual).

## Abstract

Protein O-GlcNAcylation is an evolutionary conserved post-translational modification catalysed by the nucleocytoplasmic O-GlcNAc transferase (OGT) and reversed by O-GlcNAcase (OGA). How site-specific O-GlcNAcylation modulates a diverse range of cellular processes is largely unknown. A limiting factor in studying this is the lack of accessible techniques capable of producing homogeneously O-GlcNAcylated proteins, in high yield, for *in vitro* studies. Here, we exploit the tolerance of OGT for cysteine instead of serine, combined with a co-expressed OGA to achieve site-specific, highly homogeneous mono-glycosylation. Applying this to DDX3X, TAB1, and CK2α, we demonstrate that near-homogeneous mono-S-GlcNAcylation of these proteins promotes DDX3X and CK2α solubility and enables production of mono-S-GlcNAcylated TAB1 crystals, albeit with limited diffraction. Taken together, this work provides a new approach for functional dissection of protein O-GlcNAcylation.

## Introduction

O-GlcNAcylation, the decoration of serine and threonine residues on nucleocytoplasmic proteins with a single moiety of N-acetylglucosamine (GlcNAc) (Torres and Hart 1984, Holt and Hart 1986), has roles in the regulation of signal transduction (Pathak, Borodkin et al. 2012), transcription (Lewis, Burlingame et al. 2016), translation (Jang, Kim et al. 2015), and cellular viability among others (Shafi, Iyer et al. 2000, Zhu, Liu et al. 2015, Lee, Kim et al. 2020, Zhu, Cheng et al. 2020). Only two enzymes are known to be responsible for the dynamics of O-GlcNAcylation in eukaryotes, diverging from other PTMs such as phosphorylation which relies on hundreds of kinases and phosphatases (Manning, Whyte et al. 2002). A single O-GlcNAc transferase (OGT) attaches O-GlcNAc (Martinez-Fleites, He et al. 2010), and a single O-GlcNAc hydrolase (OGA) hydrolyses O-GlcNAc (Dong and Hart 1994). Unlike many other eukaryotic glycosyltransferases that have well-defined acceptor sequences and substrate selectivity’s, OGT is a more promiscuous enzyme, as the most preferred OGT recognition sequon - [TS][PT][VT]-**S/T**-[RLV][ASY] - is only found in a fraction of the reported O-GlcNAc sites in large scale O-GlcNAcomics studies (Pathak, Alonso et al. 2015, Woo, Lund et al. 2018). Indeed, mass spectrometry studies have demonstrated that OGT decorates whole regions of proteins such as the nucleoporins, HCF1, and ankyrin G in an apparently non-specific way (Trinidad, Barkan et al. 2012, Woo, Lund et al. 2018). Overall, O-GlcNAc is a widespread PTM, present on > 5,000 proteins, playing key roles in multiple cellular and physiological processes, although the site-specific functions of O-GlcNAc are poorly understood, with only a small proportion of O-GlcNAc sites functionally interrogated to date (Nie, Ju et al. 2020, Zhu, Cheng et al. 2020, Wulff-Fuentes, Berendt et al. 2021, Zhu, Zhou et al. 2022).

A key limitation in dissecting the functional consequences of site-specific O-GlcNAcylation is the difficulty in producing sufficient quantities of stoichiometrically mono-O-GlcNAcylated proteins at specific sites of interest, for use in downstream experiments. Chemical methods, such as post-translational chemical mutagenesis using dehydroalanine (dha) as a reactive handle for subsequent attachment of PTM mimics, have been used previously to generate homogenous, mono-GlcNAcylated protein mimetics (Chalker, Lercher et al. 2012, Dadová, Galan et al. 2018). However intrinsic drawbacks in using dha-chemistry to produce protein species carrying PTMs include the uncontrolled production of a racemic mixture of D/L enantiomers at the Cα, which may explain reduced recognition of the modified protein by endogenous PTM processing enzymes. For example, acetylated histones are poorly recognised by histone deacetylases after dha-chemistry (Chalker, Lercher et al. 2012). Furthermore, dha-derived O-GlcNAcylated proteins have an extra methylene linker in the side chain that is absent in native proteins (Chalker, Lercher et al. 2012, Dadová, Galan et al. 2018). Other methods such as expressed protein ligation of semi-synthetic O-GlcNAcylated proteins (Muir, Sondhi et al. 1998), have been used to investigate the role of O-GlcNAc on α-synuclein (Marotta, Lin et al. 2015), and HSP27 (Balana, Levine et al. 2021). However, this requires a desulfurization step that converts all native cysteines to alanine (Balana, Levine et al. 2021), necessitating extensive control experiments to ensure that global desulfurization does not affect the protein folding or function.

The use of bacterial systems to co-express OGT with recombinant protein substrates has also been explored (Shen, Gloster et al. 2012, Gao, Shi et al. 2018, Li, Yang et al. 2023). This approach takes advantage of the lack of OGT in the *E. coli* genome and the absence of bacterial OGT substrates (Lubas, Frank et al. 1997). Therefore, only the desired substrate is GlcNAc modified in bacterial expression systems (Gao, Shi et al. 2018). However, while some substrates, like TAB1 or histone H2B, produce high yields of soluble O-GlcNAc modified protein (Shen, Gloster et al. 2012, Gao, Shi et al. 2018), others, like the crumbs cell polarity complex 1 (CRB1) or the Abelson tyrosine-Kinase 2 (ABL2-FABD), become insoluble following co-expression with OGT (Goodwin, Thomasson et al. 2013). Furthermore, it has been shown that over-expression of OGT and substrates in bacterial systems can lead to modification of off-target sites, which may not reflect the physiological O-GlcNAcylation patterns observed in eukaryotes (Fig.1A) (Shen, Gloster et al. 2012, Goodwin, Thomasson et al. 2013). The lack of straightforward and accessible methods for producing stoichiometric and homogeneous quantities of O-GlcNAcylated proteins at specific sites, in high yields, remains as one of the main technical hurdles in the O-GlcNAc field.

**Figure 1.**
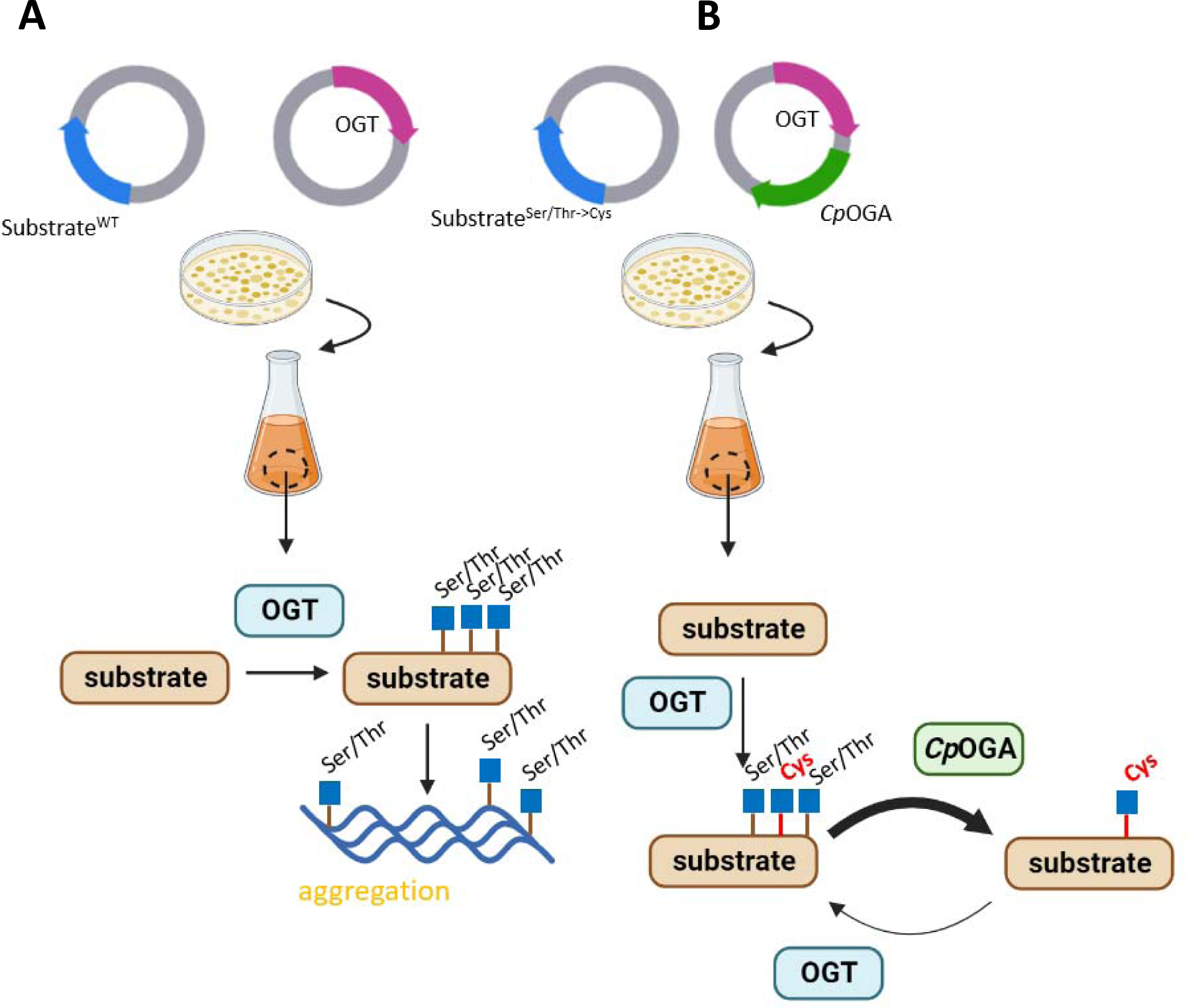
Illustrated summary of the dual cysteine mutagenesis/ *in cellulo* O-GlcNAc stripping approach to producing highly homogenous mono-S-GlcNAcylated proteins. A) Previously, OGT substrates of interest were co-expressed with OGT in *E. coli*. This process introduces multiple O-GlcNAc sites and produces a heterogenous sample of variable yield and purity. B) The method reported in this study, mutating the O-GlcNAc site of interest from Ser/Thr to Cys followed by *in cellulo* stripping of O-GlcNAc sites by *Clostridium perfringens* OGA, is illustrated.

Recently, we reported a genetic recoding approach to dissect site-specific O-GlcNAcylation (Gorelik, Bartual et al. 2019). We demonstrated that mutation of endogenous O-GlcNAc sites to Cys leads to isosteric S-GlcNAcylation within an otherwise unaltered O-GlcNAcome (Gorelik, Bartual et al. 2019). *In cellulo* experiments in eukaryotic cells with the S-GlcNAc mimetic demonstrated that OGA is unable to hydrolyse S-GlcNAc, resulting in elevated GlcNAcylation stoichiometries on OGA^S405C^, TAB1^S395C^, and FEN1^S352C^ (Gorelik, Bartual et al. 2019, Tian, Zhu et al. 2021). Here, we exploit this approach in a recombinant system to produce highly homogenous, site-specifically and near-stoichiometrically mono-S-GlcNAcylated recombinant proteins. We demonstrate that bacterial co-expression of OGT with a highly active OGA from *Clostridium perfringens* and OGT substrates carrying cysteine mutants at desired O-GlcNAc sites produces mg quantities of near homogenous mono-S-GlcNAcylated protein. Applying this approach to TAB1, casein kinase 2α (CK2α), and DEAD box RNA helicase 3 X-linked (DDX3X), we demonstrate that this rescued the solubility of DDX3X and CK2α which otherwise precipitate when only co-expressed with OGT, indicating that this approach may be broadly applicable to a range of OGT substrates and target sites. Moreover, applying this approach enabled crystallisation of mono-S-GlcNAcylated TAB1, whereas poly-GlcNAcylated TAB1 precipitated as observed previously. Therefore, this method may facilitate interrogation of the effects of O-GlcNAcylation on protein structure and function.

## Results and Discussion

### Co-expression of OGT and TAB1 produces heterogeneous O-GlcNAcylation

We previously published stable complexes of OGT with peptide substrates by fusing a truncated version of OGT to peptides containing known GlcNAc sites (Rafie, Raimi et al. 2017). This method, initially intended for the structural determination of the canonical OGT recognition sequon, showed the presence of off-target GlcNAc modification (Fig. 2A) (Rafie, Raimi et al. 2017). To determine whether this off-target activity is limited to peptide substrates, we transformed BL21 (DE3) bacterial cells already containing an OGT expressing plasmid with a His-tagged human TAB1 construct (hTAB1). hTAB1 was purified to high yield and the extent of hTAB1 O-GlcNAcylation assessed by conjugating GlcNAc moieties to GalNAc azide (GalNAz), using the azide group as a handle to attach a polyethyleneglycol (PEG) mass tag (Darabedian, Thompson et al. 2018). Using this PEGylation technique, we observed a heterogenous GlcNAc profile for the recombinant hTAB1 with up to four different GlcNAc moieties per molecule of hTAB1 protein (Fig. 2B). However, it is widely accepted that hTAB1 contains a single functional GlcNAc site on Ser395, which has been validated both by mass spectrometry and functional assays (Pathak, Borodkin et al. 2012, Woo, Lund et al. 2018, Joiner, Hammel et al. 2021). Taken together, these data show that co-expression of OGT and TAB1 produces heterogeneous O-GlcNAcylation.

**Figure 2.**
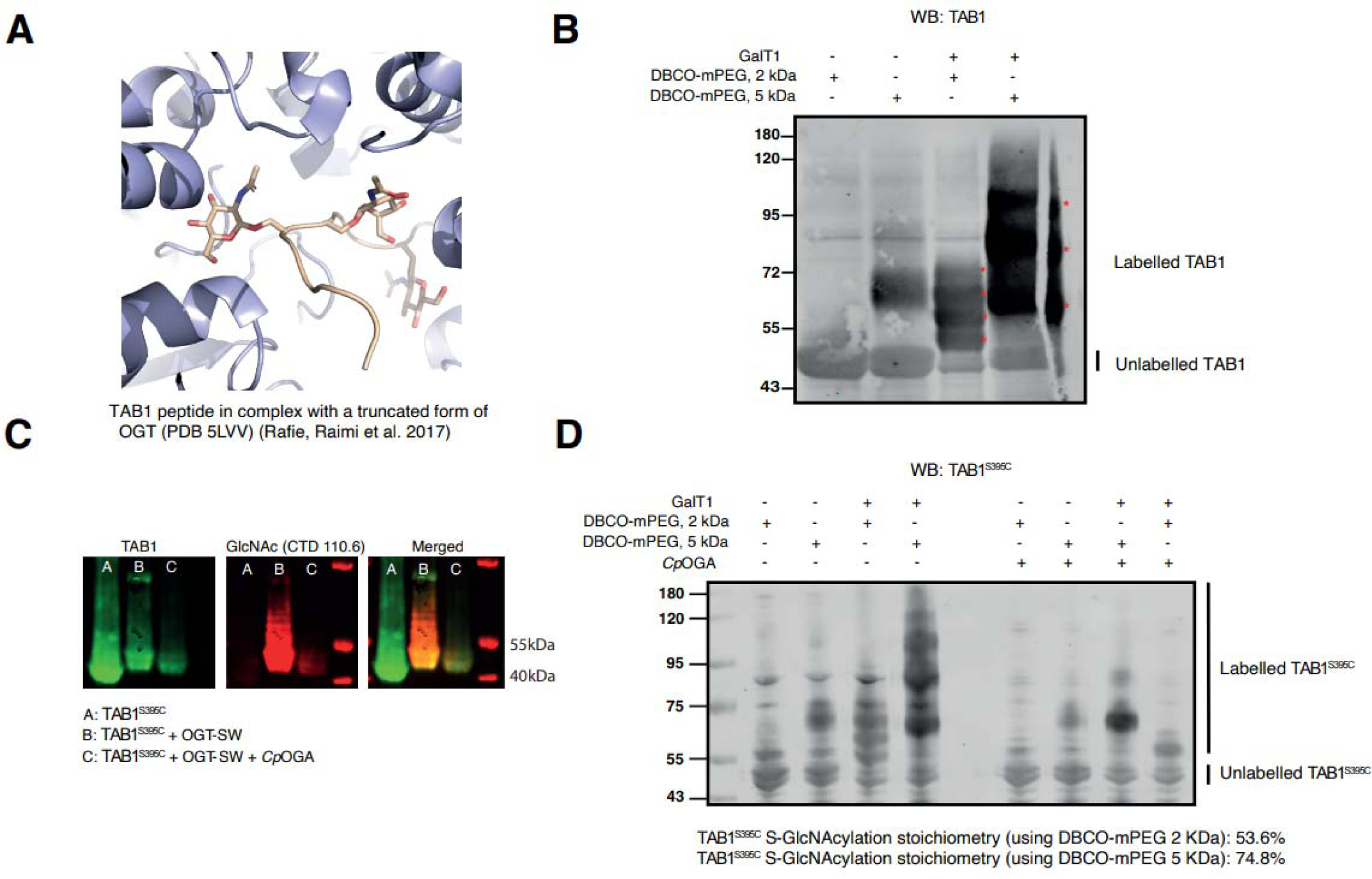
Tab1^S395C^ co-expression with OGT/ *Cp*OGA removes off-target O-GlcNAcylation sites and yields a near stoichiometrically modified S-GlcNAc protein. a) Crystal structure of a TAB1 peptide, including the canonical Ser395 O-GlcNAc site, fused to OGT shows off-target GlcNAcylation (PDB: 5LVV) b) 1 μg of purified Tab1 co-expressed with OGT was azide labelled with GalT1 and conjugated to either DBCO-PEG (2 kDa) or DBCO-PEG (5 kDa) via strain promoted azide-alkyne cycloaddition. Mass shifts compatible with the presence of several GlcNAc moieties were detected by Western blot against TAB1. c) Confirmation of TAB1^S395C^ GlcNAcylation by Western blot using anti-O-GlcNAc CTD 110.6 antibody (red channel). Lane A contains TAB1^S395C^, lane B contains TAB1^S395C^ co-expressed with OGT and lane C contains TAB1^S395C^ co-expressed with OGT and *Cp*OGA. The identity of TAB1 was confirmed by using an anti-TAB1 antibody (green channel). d) 1 μg of purified Tab1^S395C^ was azide labelled with GalT1 and conjugated to either DBCO-PEG (2 kDa) or DBCO-PEG (5 kDa) via strain promoted azide-alkyne cycloaddition. PEGylated glycoproteins were detected via Western blotting for Tab1.

### A bacterial system to produce recombinant mono-GlcNAcylated TAB1

Off-target O-GlcNAcylation (i.e., O-GlcNAcylation occurring at non-physiological O-GlcNAc sites) may alter protein activity, stability, or protein:protein interactions, complicating interpretation of downstream *in vitro* studies. Additionally, different O-GlcNAcylation sites may exert different regulatory effects on proteins, challenging interpretation of *in vitro* studies using heterogenous poly-O-GlcNAc proteins. We therefore explored the possibility of permanently installing a non-cleavable, single GlcNAc moiety in OGT substrates using bacterial host expression systems ((Gorelik, Bartual et al. 2019) ; see Fig. 1B for an illustrated summary). As a first step we mutated the single reported functional TAB1 GlcNAc site Ser395 to cysteine (TAB1^S395C^) and demonstrated that this can still be modified by co-expression with OGT (Fig. 2C). However, like the wild type TAB1 co-expressed with OGT (Fig. 2B), we detected 4 GlcNAc sites on the resulting GlcNAcylated TAB1^S395C^ by PEGylation (Fig. 2D). We next took advantage of the inability of glycoside hydrolases to hydrolyse thioglycosides (De Leon, Levine et al. 2017, Gorelik, Bartual et al. 2019), by co-expressing TAB1^S395C^ with a customized plasmid harbouring both OGT and the highly active bacterial O-GlcNAcase *Cp*OGA (Rao, Dorfmueller et al. 2006), anticipating this would lead to retention of a single S-GlcNAc site, with removal of spurious O-GlcNAcylation. Indeed, co-expression with *Cp*OGA resulted in TAB1 carrying a single GlcNAc modification as assessed by the PEG mass tagging approach, although a faint band at ∼80 kDa was observed following mass tagging with DBCO-mPEG (5 kDa) which could represent residual di-glycosylation of TAB1. This additional band could also represent non-specific thiol labelling by the DBCO probe as a faint band shift is also observed in the unlabelled control without GalT1 (Fig. 2D). Nevertheless, the possibility remained that residual di-glycosylation of TAB1 occurred even after co-expression with *Cp*OGA. Therefore, to investigate possible residual sample heterogeneity and thus validate whether TAB1^Ser395Cys^ was mono-S-GlcNAcylated, we analysed both TAB1^WT^ and TAB1^S395C^ after co-expression with either OGT or both OGT and *Cp*OGA by mass spectrometry (see supplementary file 1 for list of detected peptides and supplementary Fig. S4A,B for representative spectra). After co-expression of TAB1^WT^ with OGT, a C-terminal peptide (387-VYPVSVPYSSAQSTSK-402) was detected in both the mono- and di-glycosylated state. After co-expression with both OGT and *Cp*OGA only the unmodified version of the peptide was detected, indicating successful stripping of O-GlcNAc sites by *Cp*OGA. From the sample with TAB1^S395C^ co-expressed with OGT we identified three peptides carrying one GlcNAc modification (337-IHSDTFASGGER-348, 364-NFGYPLGEMSQPTPSPAPAAGGR-386 and 387-VYPVSVPYCSAQSTSK-402). The peptide VYPVSVPYCSAQSTSK does not carry a carbamidomethyl modification from the iodoacetamide alkylation, which indicates that the modification is located at Cys395 (see supplementary file 1). In contrast to the two other peptides, the VYPVSVPYCSAQSTSK peptide was still identified as modified with GlcNAc after co-expression with both OGT and *Cp*OGA. Therefore, TAB1^S395C^ is mono-S-GlcNAcylated following co-expression with OGT and *Cp*OGA, with no detectable residual modification of off-target sites. As mentioned above, O/S-GlcNAc site mapping identified 2 and 3 sites of modification for TAB1^WT^ and TAB1^S395C^, respectively, in contrast to PEGylation data which indicated 4 sites of modification for both constructs following co-expression with OGT. This discrepancy may stem from the poor ionisation efficiency of the GlcNAc peptide or non-specific binding of the TAB1 antibody after PEGylation to contaminant proteins. Taken together, these data show that we have generated a bacterial system to generate recombinant mono-GlcNAcylated TAB1.

### Crystallization of mono-S-GlcNAcylated TAB1

Determining the atomic structure of any macromolecule requires a pure and homogeneous protein sample. Failure in obtaining complete PTM stoichiometry often leads to a decrease in the sample’s homogeneity which significantly reduces the likelihood of success (McPherson and Gavira 2014). It is worth noting that there are currently no experimentally determined protein structures with structurally defined O-GlcNAc sites. Indeed, we have previously unsuccessfully attempted to crystallise GlcNAcylated TAB1. Failure to achieve this is probably linked to the TAB1 heterogeneity when co-expressed with OGT (Fig. 2B). We wondered if our mono-GlcNAcylated TAB1^S395C^ would improve the chances of crystallising GlcNAcylated TAB1. Under our experimental conditions, TAB1^S395C^ purified from the co-expressed system with OGT and *Cp*OGA produced well-defined hexagonal crystals while TAB1^S395C^ co-expressed only with OGT failed to crystallise and instead precipitated (Supplementary Fig. 1A). A non-modified TAB1^S395C^, used as an experimental control, produced the same hexagonal-shaped crystals as the mono-GlcNAcylated TAB1^S395C^. Further analysis of the TAB1^S395C^ crystals’ content by Western blot ruled out the possibility of the crystals being formed by non-modified protein (Supplementary Fig. 1B). However, these crystals only diffracted to low resolution and did not allow structure solution. Additional analysis by differential scanning fluorimetry showed no differences in the thermal stability of S-GlcNAcylated TAB1^S395C^ versus un-modified TAB1^S395C^ (Supplementary Fig. 1C). Moreover, poly-GlcNAcylated TAB1, produced by co-expression with OGT only, also showed no differences in thermal stability. Poly-GlcNAcylated TAB1, produced by co-expression with full-length OGT in *E. coli* with up to 5 sites of O-GlcNAcylation, was protected against thermally-induced aggregation in a previous study (Yuzwa, Shan et al. 2012). Our inability to detect a similar thermoprotective effect of GlcNAc on poly-GlcNAcylated TAB1 likely stems from using a shorter OGT construct lacking the 9 most N-terminal TPRs, as opposed to full-length OGT, which may explain the resulting modification of fewer off-target sites compared to the previous study. These data therefore highlight the importance of studying site-specific effects *in vitro* using the S-GlcNAc approach when assigning functionality to O-GlcNAcylation events. Overall, to the best of our knowledge these data demonstrate the first successful crystallisation of a GlcNAcylated protein – TAB1 - facilitated by the approach of generating stoichiometrically mono-S-GlcNAcylated proteins.

### Mono-S-GlcNAcylation of DDX3X and CK2α rescues protein solubility

For some OGT substrates, previous attempts to produce soluble recombinant GlcNAc modified protein were unsuccessful, owing to the O-GlcNAcylated form of the protein being insoluble (Goodwin, Thomasson et al. 2013), possibly due to off-target GlcNAcylation that might either affect solubility or disrupt protein folding. We next explored whether restriction to homogeneous mono-S-GlcNAcylation of physiologically relevant sites would remove these issues. An example of this phenomenon is DEAD box RNA helicase 3 X-linked (DDX3X), an RNA helicase with diverse roles in transcriptional regulation and translation initiation (Lai, Chang et al. 2010, Chen, Yu et al. 2016, Cannizzaro, Bannister et al. 2018). A previous study linked DDX3X Ser584 O-GlcNAcylation to OGT intellectual disability aetiology (Mitchell, Czajewski et al. 2022). *In vitro* dissection of the role of O-GlcNAc on DDX3X is hampered by the aggregation of DDX3X following co-expression with OGT (Fig. 3A, lanes 4-6). Indeed, for *E. coli* expression a soluble construct of DDX3X was used (residues 132-607) that has been crystallised previously (Floor, Condon et al. 2016). Despite this, the O-GlcNAcylated form of DDX3X accumulated in the insoluble pellet fraction (Fig. 3A, lane 5), and we were unable to detect soluble DDX3X protein by Western blot after one round of Ni^2+^-affinity purification (Fig. 3A, lane 6). Furthermore, co-expression of DDX3X with OGT and *Cp*OGA rescued yields of soluble DDX3X (Fig. 3A, lanes 7-9), suggesting off-target O-GlcNAcylation may impair the folding and/ or solubility of DDX3X. DDX3X^S584C^ was therefore co-expressed with OGT and *Cp*OGA, to investigate whether mono-S-GlcNAcylation of DDX3X would rescue protein solubility. GlcNAcylated DDX3X^S395C^ was successfully purified from *E. coli* lysate, as indicated by Western blotting, in high yield (average yield across three independent experiments: ∼ 1.25 mg per L of *E. coli* culture), and no GlcNAcylated DDX3X^S395C^ was observed in the pellet fraction (Fig. 3A, lanes 13-15). This result was in sharp contrast to DDX3X^S395C^ co-expressed with OGT only, which similarly to DDX3X^WT^ showed accumulation of GlcNAcylated DDX3X^S584C^ in the insoluble fraction and reduced yield of soluble protein (Fig. 3A, lanes 10-12). Therefore, targeted site-specific S-GlcNAcylation of DDX3X rescued solubility of the GlcNAcylated form of the protein.

**Figure 3.**
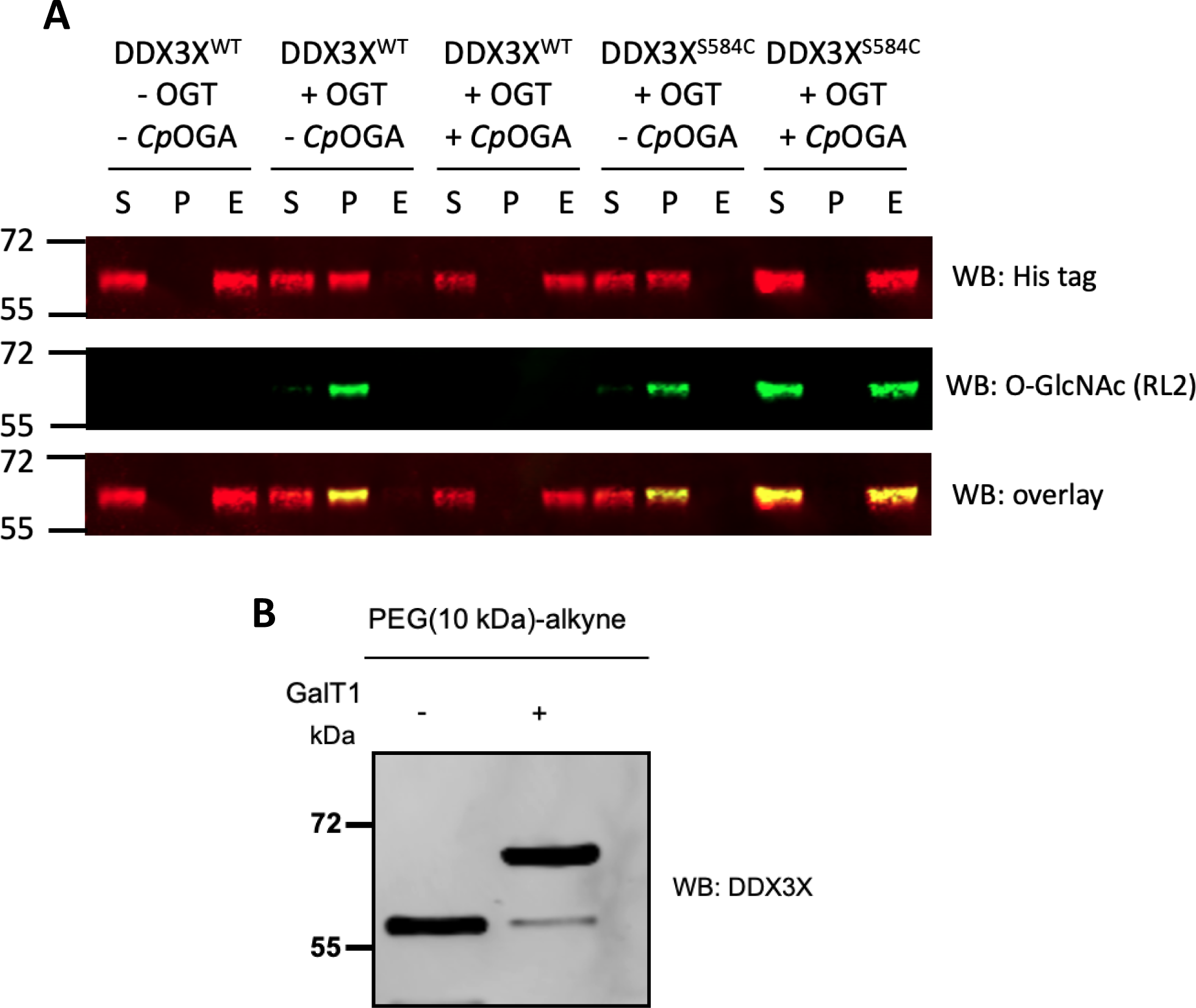
DDX3X^S584C^ is near homogenously mono-S-GlcNAcylated at C584 following OGT/*Cp*OGA co-expression. a) His_6_-DDX3X^WT^ His_6_-DDX3X^S584C^ (residues 132-607) was co-expressed with either OGT only or OGT and *Cp*OGA in *E. coli*. After one round of Ni^2+^-affinity purification, the soluble fraction (S), insoluble pellet fraction (P), and imidazole elution following Ni^2+^-affinity purification were analysed for the presence of the N-terminal His tag and O-GlcNAcylation by WB. b) Copper catalysed azide alkyne cycloaddition of DDX3X^S584C^, after co-expression with OGT and *Cp*OGA, to alkyne-PEG (10 KDa). A separate labelling reaction without GalT1 was carried out as a negative control.

The stoichiometry of S-GlcNAc modification on DDX3X^S395C^ was subsequently investigated by PEGylation of DDX3X^S395C^ GlcNAc sites to alkyne-PEG (10 kDa). A single band shift was observed which, upon densitometric analysis, revealed a stoichiometry of 95% mono-S-GlcNAcylation (Fig. 3B). Furthermore, as mutagenesis of Ser584 has not previously been reported, we verified by DSF that mutagenesis of Ser584 to Cys does not affect the thermal stability and, by extension, the folding of DDX3X (Supplementary Fig. 2). Therefore, cysteine mutagenesis of Ser584 on DDX3X, paired with OGT/*Cp*OGA co-expression, enables expression and purification of near-homogenous mono-S-GlcNAcylated DDX3X at a targeted site, in high yield, for subsequent *in vitro* studies.

To assess the broader applicability of this approach, we next targeted the known GlcNAc site (Ser347) on casein kinase 2α (CK2α) (Tarrant, Rho et al. 2012). CK2α is dual specificity Ser/Thr kinase that forms part of the CK2 holoenzyme, along with CK2β (Niefind, Guerra et al. 2001), and plays essential roles in cell survival (Yamane and Kinsella 2005), DNA replication (Feng, Lu et al. 2021), and development (Götz and Montenarh 2017). O-GlcNAcylation of a single site – Ser347 – destabilises CK2α (Tarrant, Rho et al. 2012), and modulates CK2 kinase activity (Schwein, Ge et al. 2022). In the same fashion as TAB1 and DDX3X, we co-expressed CK2α^WT^ and the cysteine mutant (CK2α^S347C^) with OGT +/-*Cp*OGA. We observed that co-expression of both CK2α^WT^ and CK2α^S347C^ with OGT alone resulted in aggregation of >50% of the O/S-GlcNAcylated glycoform of the protein (Fig 4A, lanes 4-6, 10-12). This was reflected in the poor yields of soluble CK2α^WT^ and CK2α^S347C^ following co-expression with OGT, compared to the yields of CK2α^WT^ and CK2α^S347C^ expressed without OGT (Supplementary Fig. 3). However, co-expression of CK2α^WT^ and CK2α^S347C^ with both OGT and *Cp*OGA improved soluble protein yields across two independent experiments, suggesting that hyper-O-GlcNAcylation of off-target CK2α O-GlcNAc sites causes protein aggregation in *E. coli*. We additionally observed by O-GlcNAc Western blot that CK2α^WT^was not GlcNAcylated following co-expression with OGT/*Cp*OGA, whereas CK2α^S347C^ was (Fig 4A). Utilising the PEG mass tagging approach, we determined that CK2α^S347C^ was mono-S-GlcNAcylated at a stoichiometry of >95% (Fig. 4B). Although a putative di-glycosylation signal can be observed at ∼75 kDa, the absence of this band in the unlabelled control without GalT1 and its simultaneous appearance following PEGylation of un-modified CK2α (Fig. 4B, lane 2) suggests this band results from non-specific binding of the His tag antibody to bacterial protein contaminants in the commercial GalT1 stock, a phenomenon reported previously (Darabedian, Thompson et al. 2018).Taken together these data show that the bacterial OGT/*Cp*OGA co-expression system combined with targeted Ser->Cys mutations leads to mono-S-GlcNAcylation of DDX3X and CK2α and increased solubility.

**Figure 4.**
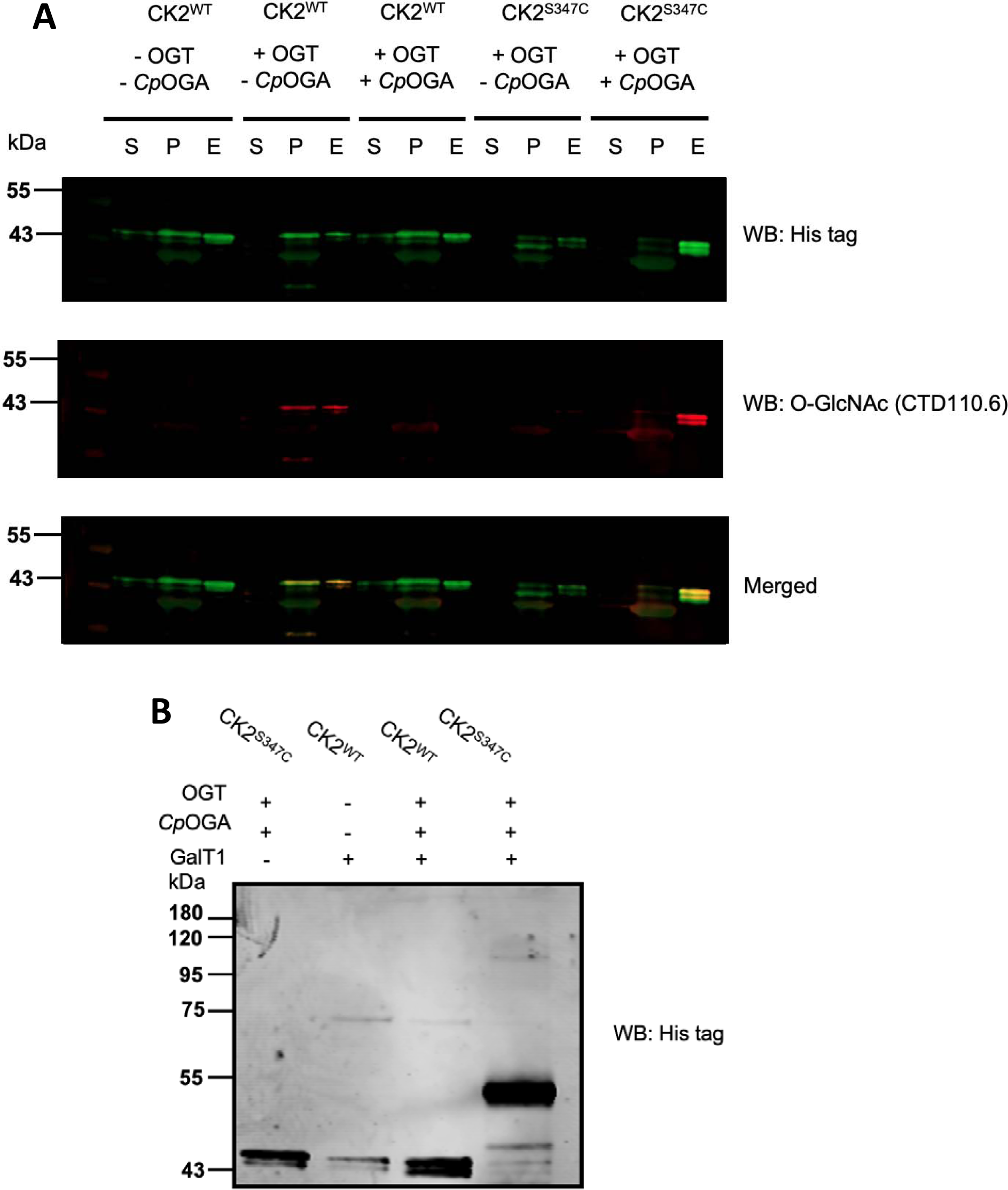
S-GlcNAcylation of CK2α^S347C^ after OGT/*Cp*OGA co-expression yields stoichiometrically mono-S-GlcNAcylated CK2α and rescues protein solubility. a) *E. coli* were transformed with plasmids encoding either CK2α only, CK2α with OGT, or CK2α with OGT/*Cp*OGA. After induction with IPTG, soluble (S) and insoluble fractions (P) of *E. coli* lysate, and the elution from Ni^2+^ beads following Ni^2+^-affinity chromatography (E), were analysed by blotting for the N-terminal His tag and O-GlcNAc. b) O/S-GlcNAc sites on purified CK2α^WT^, expressed in the presence or absence or OGT/*Cp*OGA, and CK2α^S347C^, expressed with OGT/*Cp*OGA, were azide-labelled via GalT1 and conjugated to DBCO-mPEG (5 kDa) via strain promoted azide-alkyne cycloaddition. Total CK2α (both labelled and unlabelled) was detected by blotting for the N-terminal His tag. CK2^S347C^ was reacted with DBCO-mPEG (5 kDa) in the absence of GalT1 azide labelling to control for non-specific labelling of cysteine thiols by the DBCO probe.

## Discussion

O-GlcNAcylation modulates diverse cellular and developmental processes, from stem cell pluripotency through to bioenergetics (Jang, Kim et al. 2012, Zhu, Zhou et al. 2022), although the role of O-GlcNAc is only understood for a small fraction of the >5,000 O-GlcNAc proteins identified to date. Elucidating the site-specific effects of O-GlcNAc on protein function will be key to understanding the myriad roles O-GlcNAc plays in cellular processes. However, the study of the site-specific effects of GlcNAc on particular proteins has historically proven challenging. Progress in the O-GlcNAc field towards site-specific installation of O-GlcNAc or GlcNAc mimetics has lagged behind compared to other PTMs, such as phosphorylation, where Ser -> Asp/Glu mutagenesis has been routinely implemented to mimic and study specific phosphorylation events. The study, especially at the molecular level, of the effects of O-GlcNAcylation requires accessible, and scalable tools for producing mg quantities of stoichiometrically GlcNAcylated proteins, modified at a single target site of interest.

Co-expression of OGT with protein substrates has previously been used to study the effects of O-GlcNAc on Tau and malate dehydrogenase 1, among other targets (Gao, Shi et al. 2018, Zhu, Zhou et al. 2022, Li, Yang et al. 2023). This method provides a considerable simplicity over the most demanding and comparatively expensive chemical alternatives only affordable by a handful of very specialised laboratories worldwide (Dadová, Galan et al. 2018, Balana, Levine et al. 2021). However, this approach is not without issues as several groups, including ourselves, have observed that co-expression with OGT often leads to off-target modifications which could lead to undesired effects. (Goodwin, Thomasson et al. 2013, Rafie, Raimi et al. 2017). This, together with the possibility of the bacterial NagZ hydrolase acting as a putative GlcNAc hydrolase (Goodwin, Thomasson et al. 2013), potentially complicates data derived from proteins produced by co-expression with OGT.

We have previously demonstrated that OGT is capable of S-GlcNAcylating Cys thiol groups *in cellulo* after mutagenesis of Ser/Thr O-GlcNAc sites to Cys, while the endogenous OGA is unable to hydrolyse the S-GlcNAc moiety (Gorelik, Bartual et al. 2019). Therefore, to compensate for the spurious off-target O-GlcNAcylation observed in the traditional OGT co-expression approach, we mutated the Ser/Thr O-GlcNAc sites on several target proteins to Cys. Then, to remove any off-target O-GlcNAc sites, we co-expressed OGT and substrates with the highly efficient *Clostridium perfringens* OGA (Rao, Dorfmueller et al. 2006). As we observed in all our tested proteins, this approach leaves a single S-GlcNAc moiety attached to the OGT substrate and leads to the observed site-specific elevation of GlcNAcylation stoichiometries. The ability to study the effects of individual GlcNAcylation events may be useful in instances where multiple O-GlcNAcylation sites on the same protein exert different effects, e.g. on protein activity and localisation. For example, O-GlcNAcylation of SP1 at Ser491 blocks its interaction with transcription initiation factor (TAF) 110D (Roos, Su et al. 1997), whereas modification of Thr421, Ser613, Ser699, and Ser703 may regulate SP1 localisation in response to insulin signalling (Majumdar, Harrington et al. 2006). Therefore, studies of multi-site O-GlcNAcylation, which require differentiating the effects of different O-GlcNAcylation sites on protein function, may be facilitated by the approach we describe here.

Interestingly, we observed that co-expression of DDX3X or CK2α with OGT resulted in precipitation of the O-GlcNAcylated glycoform of the protein. A similar phenomenon has been reported previously for CRB1 and ABL2, where the O-GlcNAcylated glycoforms of either protein was exclusively found in the insoluble fraction – as we observed for full length DDX3X co-expressed with OGT (Goodwin, Thomasson et al. 2013). This may stem from co-translational O-GlcNAcylation of OGT substrates (Zhu, Liu et al. 2015), which could interfere with the protein surface electrostatic properties or prevent proper folding. Additionally, NagZ in *E. coli* may reduce stoichiometry by stripping GlcNAc, an issue which cannot be overcome by treating *E. coli* with the OGA inhibitor PUGNAc (Haltiwanger, Grove et al. 1998, Goodwin, Thomasson et al. 2013). Indeed, a recent study utilising OGT co-expression with malate dehydrogenase 1 yielded only 31% stoichiometry, which may at least partially be attributable to NagZ activity (Zhu, Zhou et al. 2022). These issues previously limited the utility of *E. coli* for producing large quantities of GlcNAcylated protein for study. However, the ability to recombinantly produce hydrolase resistant mono-S-GlcNAcylated TAB1, DDX3X and CK2α using our approach suggests that these limitations can now be overcome.

The ability to purify mg quantities of recombinant mono-S-GlcNAcylated proteins opens many avenues for future research. For example, similar to the effect observed in a TAB1 peptide-OGT fusion construct (Rafie, Raimi et al. 2017) and replicated again here with the isolated TAB1 protein, the spurious GlcNAcylation of TAB1 by OGT produces a complex, heterogeneously GlcNAcylated, sample. Furthermore, it is possible that the very same effect could be one of the reasons behind the lack of an O-GlcNAcylated protein in structural databases. However, as the samples produced by our approach are near-stoichiometrically mono-S-GlcNAcylated, these samples are highly homogenous and thus more amenable to crystallography as demonstrated by the successful crystallisation of mono-S-GlcNAcylated TAB1^S395C^ reported here. Obtaining the atomic-resolution structure of GlcNAcylated proteins is of the utmost importance as these may provide insights into the effects of GlcNAcylation on protein folding, protein-protein interactions, or protein conformation, the latter permitting study of structure-activity relationships. This is particularly relevant considering recent studies demonstrating, by thermal profiling, that O-GlcNAc can act as both a stabiliser and destabiliser of different proteins, a phenomenon that could be explained by GlcNAc-dependent conformational changes in the protein (King, Serrano-Negrón et al. 2022).

Substitution of Ser/Thr for Cys can be achieved by conventional molecular cloning techniques, and production of mono-S-GlcNAcylated proteins requires a basic set-up for *E. coli* culture and protein purification that is widely available across many research groups. Due to the relatively recent discovery that OGT tolerates Cys to produce S-GlcNAc, this approach needs further validation on a greater number of OGT substrates, especially if used in *in vivo* biological systems. Nevertheless S-GlcNAc so far presents potentially the most straight forward and effective approach to study the effects of protein mono-GlcNAcylation both *in vivo* and especially in *in vitro* experimental setups.

## Materials & Methods

### Cloning of constructs for E. coli expression

The region of DDX3X 132-607 was ordered as a codon optimised geneblock from Integrated DNA Technologies including an N-terminal *Bam*HI site, a C-terminal stop codon and a *Not*I site. This fragment was digested and cloned into a *Bam*HI-*Not*I digested and dephosphorylated vector pHEX6P1 (this proprietary vector is based on pGEX6P1 with a 6xHis tag in place of the GST tag). The CK2α (1-361) and TAB1 (7-410) inserts were cloned in the same way into the same vector. All inserts were confirmed by DNA sequencing. The mutations in DDX3X (Ser584Ala and Ser584Cys), CK2α (Ser347Cys) and TAB1 (Ser395Cys) were carried out by site directed mutagenesis and verified by DNA sequencing.

### Expression and purification of OGT substrates

A list of the proteins used in this work, with associated tags and construct boundaries, can be found in Supplementary Table 1. Briefly, each protein construct was transformed in either BL21 DE3 pLysS (NEB) harbouring a truncated OGT version with an intact catalytic core (OGT^SW^) (Lazarus, Nam et al. 2011) or transformed in BL21 DE3 pLysS cells already containing a co-expression plasmid which simultaneously expresses tagless OGT^SW^ and tagless *Cp*OGA. Each protein was expressed by inoculating 5 mL of overnight culture into 1 L of LB broth supplemented with ampicillin and kanamycin (total volume: 6 L). At an OD of 0.6, cultures were induced with the addition of 250 μM IPGT (Formedium) and grown at 16-18 4 °C overnight (except for CK2α, see below). Cultures were then recovered by centrifugation (4,000 x g, 4 °C, 40 min in a Beckman Avanti JXN-26 (rotor: JLA 8.1)) and pellets resuspended in approximatively 30 mL of lysis buffer (20 mM HEPES, pH 7.5, 250 mM NaCl, 30 mM imidazole, 0.5 mM TCEP, 1 mM benzamidine, 0.2 mM PMSF, 5 μM leupeptin). Samples were subsequently stored at −20 °C until further use.

For expression of CK2α, a previously described protocol (Olsen, Boldyreff et al. 2006) was used. Briefly, *E. coli* expressing the plasmid encoding CK2α were grown to an OD of 0.5, followed by induction with 300 μM IPTG for 2 h (37 °C). Cells were harvested and lysed as described above in lysis buffer (20 mM Tris-HCl, pH 7.4, 200 mM NaCl, 0.5 mM TCEP, supplemented with 1 mM benzamidine, 0.2 mM PMSF, 5 μM leupeptin). All downstream purification steps were identical to those detailed below.

Frozen samples were defrosted and supplemented with ∼0.1 mg/mL lysozyme, and ∼0.1 mg/mL bovine DNase (Sigma Aldrich). Sample lysis was achieved with a French press pressure cell homogeniser adjusted to 1500 psi (Thermo Fisher Scientific). Then, lysates were centrifuged at 60,000 x g (4 °C) for 45 min (Beckman Avanti JXN-26 (rotor: JA 25.5)) and immediately supplemented with 10 mL pre-washed Ni^2+^-NTA agarose (QIAGEN) followed by incubation for 2 h at 4 °C with rotation. Beads containing the protein of interest were washed twice with lysis buffer, twice with a buffer supplemented with 20 mM imidazole, to remove non-specific binders, and proteins subsequently eluted with 250 mM imidazole. Due to the nature of DDX3X as a transcription factor, one additional step of washing with 1M NaCl to remove bound nucleic acids followed by a wash with lysis buffer was introduced for this protein prior to the wash with 20 mM imidazole (Floor, Condon et al. 2016). Eluted proteins were supplemented with 150 μL 4 mg/mL recombinant PreScission protease (Leong 1999) and simultaneously dialysed overnight to cleave the 6xHis tag and remove the excess of imidazole. After dialysis, soluble protein was incubated with fresh Ni^2+^-NTA beads, and eluent containing the tagless protein of interest was then concentrated to 5 mL and loaded on a HiLoad^TM^ 16/600 superdex^TM^ 200 pg size exclusion column (GE) equilibrated with lysis buffer. Fractions containing the protein of interest at the expected Mw were pooled, buffer exchanged into 20 mM HEPES, pH 7.5, 250 mM NaCl, 0.5 mM TCEP, concentrated to 5 - 10 mg/mL and flash frozen in liquid nitrogen prior to storage at −80 °C until further use. In the case of DDX3X, as previously reported, the buffer was exchanged to 20 mM HEPES, pH 7.5, 250 mM NaCl, 500 mM NaCl, 10% (v/v) glycerol, 0.5 mM TCEP prior to storage (Floor, Condon et al. 2016).

### GalNAc-azide labelling and CuAAC PEGylation of O-GlcNAc sites

20 μg of purified protein was chloroform/methanol precipitated by sequential addition of 600 μL methanol, 150 μL chloroform, and 400 μL water, followed by centrifugation. After removal of the upper (aqueous) phase, 450 μL MeOH was added, and protein pelleted by centrifugation. Proteins were re-suspended in 20 μL of 20 mM HEPES, pH 7.9, 1% (w/v) SDS, and azide-labelled using the Click-iT^TM^ O-GlcNAc Enzymatic Labelling System (Thermo Fisher Scientific). Briefly, 24.5 µL miliQ water, 40 µL labelling buffer, 5.5 µL 100 mM MnCl_2_, 10 µL 2 mM UDP-GalNAz, and 4 µL galactosyltransferase T1 Y289L were added to the 20 µL resuspension. The azide-labelling reaction was incubated at 4 °C overnight with rotation. After overnight labelling, cysteine residues were alkylated via incubation with 22.5 mM iodoacetamide for 30 minutes, followed by chloroform/methanol precipitation and resuspension in 20 μL 25 mM Tris-HCl, pH 8.0, 1% (w/v) SDS. CuAAC reactions were performed using the Click-IT Protein Reagent Buffer kit. Briefly, 20 μL of Click-IT reaction buffer, supplemented with 3.3 mM MeO-PEG-alkyne 10 kDa (final concentration), was added to the resuspended protein, followed by addition of 2 μL of 40 mM CuSO_4_, 2 μL Click-IT reaction buffer additive 1 and 4 μL Click-IT reaction buffer additive 2. CuAAC PEGylations were incubated for 2 h at RT (1300 rpm). After a final round of chloroform/ methanol precipitation, PEGylated proteins were resuspended in 45 μL 1 x LDS loading buffer for Western blotting.

### SPAAC PEGylation of GalNAc-azide labelled O-GlcNAc sites

Purified proteins were GalNAc-azide labelled as described above. For SPAAC PEGylation of azide-labelled proteins, after alkylation with 22.5 mM iodoacetamide, proteins were chloroform/ methanol precipitated and resuspended in 90 μL Tris-HCl, pH 8.0, 1% (w/v) SDS. 10 μL of 10 mM DBCO-mPEG was added to the 90 μL resuspension and incubated at 98 °C (5 min, 900 rpm). After reaction completion, PEGylated glycoproteins were chloroform/ methanol precipitated, resuspended in 45 μL 1 x LDS and analysed by Western blot.

### Western blotting

250 to 500 ng of purified recombinant protein, or 500 ng of PEGylated recombinant protein, was loaded onto a NuPAGE Bis-Tris (8% polyacrylamide) gel. After SDS-PAGE and transfer to nitrocellulose membrane, proteins were blotted using one of the following antibodies, with their respective dilutions: rabbit α-DDX3X (1:1,000 ; Proteintech), mouse α-His-tag (1:1,000 ; Proteintech), α-O-GlcNAc (RL2 ; 1:1,000; Novus Biologicals); α-O-GlcNAc (CTD110.6 ; 1:1,000 ; Cell Signalling Technology), rabbit α-Tab1 (1:1,000 ; Cell Signalling Technology), rabbit α-OGT (1:2,000 ; Sigma Aldrich).

### S-GlcNAc Tab1^S395C^ crystallisation

Recombinant TAB1^S395C^ protein was crystallised as previously described in (Conner, Kular et al. 2006). Briefly, TAB1^S395C^ protein (10mg/mL) was mixed with the mother liquor in a 1:1 drop ratio. An identical procedure was followed for TAB1^S395C^ co-expressed with OGT and TAB1^S395C^ co-expressed with OGT and *Cp*OGA. Hexagonal shaped crystals were observed after 3 to 5 days of incubation. To investigate the S-GlcNAcylation of TAB1^S395C^ crystals by Western blot, crystals were washed extensively in mother liquor prior to dissolution in lysis buffer and boiling in 1 X SDS (final concentration: 0.5 mg/mL).

### Differential scanning fluorimetry

Protein denaturing profiles were recorded with a BioRAD CFX Opus 96 Real-time PCR configured for detecting FRET signal. Briefly, 1 mg/mL of recombinant protein was mixed with SYPRO Orange at a 5X final concentration (Invitrogen), and reactions were supplemented with 50 mM HEPES, 150 mM NaCl pH 7.5 up to a final volume of 20 μL. All experiments were recorded in sextuplicate, and GraphPad Prism was used to plot and further analyse the data.

### Sample preparation prior to LC-MS/MS

After SDS-PAGE of 500 ng of purified protein, gel pieces of the expected molecular weight were excised and in-gel digested as previously described (Shevchenko, Tomas et al. 2006). Briefly, ∼1 mm x 1 mm gel pieces were treated with neat acetonitrile, followed by alkylation of free cysteine thiols with 55 mM iodoacetamide in 100 mM ammonium bicarbonate for 20 min in the dark (room temperature). Gel pieces were then destained via incubation with 1:1 (vol/vol) 100 mM ammonium bicarbonate / acetonitrile for 1 hr at room temperature. The gel pieces were saturated with 250 ng trypsin (Promega) in 50 μL 10 mM ammonium bicarbonate, pH 8.0, at 4 °C for 2 h prior to incubation for 16 h overnight at 37 °C in a thermomixer. Peptides were extracted by the addition of 100 μL 1:2 (vol/vol) 5% formic acid / acetonitrile followed by shaking for 15 min at 37 °C in a thermomixer. The extracted peptides were prepared for liquid chromatography tandem mass spectrometry (LC-MS/MS) analysis by desalting samples using Empore™ Octadecyl C18 reverse phase membranes stage tips as described in (Rappsilber, Ishihama et al. 2003).

### Mass spectrometry-based identification of O/S-GlcNAc

The desalted samples were solubilized in solvent A (0.1 % formic acid) and analysed on an Orbitrap Eclipse Tribrid Mass Spectrometer (Thermo Fisher Scientific) coupled to an Easy-nLC 1200 (Thermo Scientific) with a 2 cm trap column (100 µm I.D.) and a 15 cm analytical column (75 µm I.D.) both packed in-house with ReproSil-Pur 120 C18-AQ 1.9 µm (Dr. Maisch GmbH). Peptides were separated using a 35 min gradient from 6 % solvent B (80 % acetonitrile, 0.1 % formic acid) to 44 % solvent B at 250 nL/min. The spectral data were analysed using Proteome Discoverer™ 2.5 Software (Thermo Fisher Scientific) the Mascot Search engine (Matrix Science) with the following search parameters: trypsin digestion with ≤ 2 missed cleavages, carbamidomethylation of Cys residues, oxidation of Met residues and O-/S-GlcNAc as variable modifications. Search tolerance were 10 ppm for the precursor and 0.05 Da for the fragment ions. Data were searched against a database containing the *E. coli* proteome and both the TAB1^WT^ and TAB1^S395C^ sequence. The modification site probabilities for PSMs were determined using the IMP-ptmRS node The Mascot search results are reported with a significance threshold of p<0.005.

## Acknowledgements

This work was funded by a Wellcome Trust Investigator Award (110061), a Novo Nordisk Foundation Laureate award (NNF21OC0065969) and a research grant (VIL54496) from VILLUM FONDEN to D.M.F.v.A. C.W.M is funded by the BBSRC EASTBIO doctoral training program.

## Author contributions

D.M.F.v.A, S.G.B, and C.W.M conceived the study. A.T.F cloned all constructs used in this study. S.G.B and C.W.M performed recombinant expression of protein constructs and DSF experiments. C.W.M carried out PEGylations and stoichiometry measurements. S.G.B performed crystallisation and structural analysis. C.S. performed O/S-GlcNAc site mapping. The manuscript was written by D.M.F.v.A, S.G.B, C.W.M, and C.S.

## Abbreviations

(OGT): O-GlcNAc Transferase
(OGA): O-GlcNAc Hydrolase
(GlcNAc): N-acetyl glucosamine
(TAB1): TGFβ activated kinase 1
(CK2α): Casein Kinase 2α
(DDX3X): DEAD box RNA Helicase 3, X-linked
(HCF1): Host Cell Factor 1
(PTM): Post-Translational Modification
(dha): dehydroalanine
(CRB1): Crumbs Cell Polarity Complex 1
(ABL2): Abelson Tyrosine-Kinase 2
(*Cp*OGA): *Clostridium perfringens* OGA
(FEN1): Flap Structure-Specific Endonclease 1
(DBCO-PEG): Dibenzocycloocytne Polyethylene Glycol
(MeO-alkyne PEG): Methoxy Alkyne Polyethylene Glycol
(SPAAC): Strain Promoted Azide Alkyne Cycloaddition
(CuAAC): Copper Catalysed Azide Alkyne Cycloaddition

## Data availability statement

The data underlying this article will be shared upon reasonable request to the corresponding author.

---

**Supplementary Figure S1:**
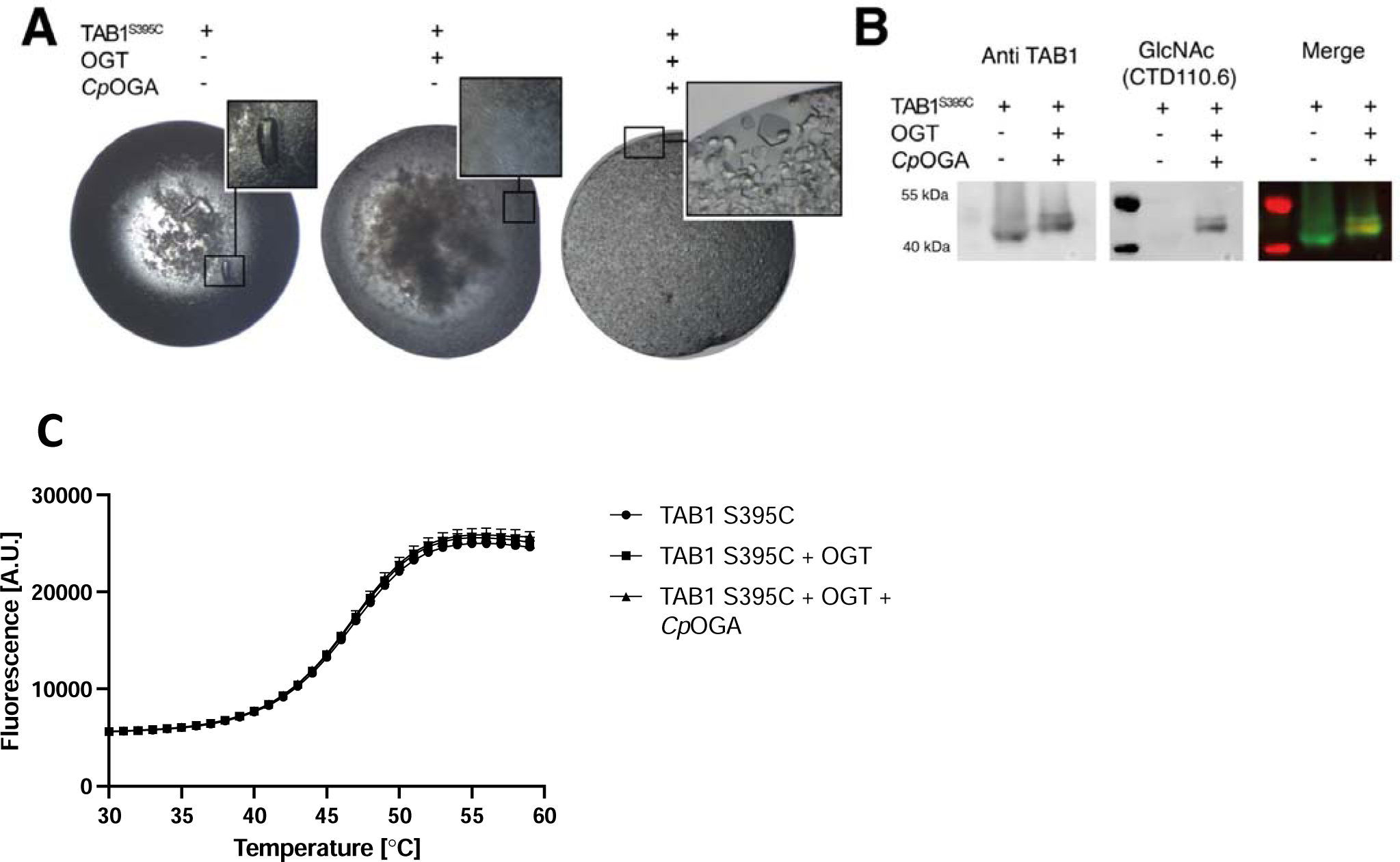
Mono S-GlcNAcylated TAB1 crystallisation and thermal stability. a) Images of the crystal trials of TAB1^S395C^ drops, from left to right: TAB1^S395C^, TAB1^S395C^ co-expressed with OGT and finally, TAB1^S395C^ co-expressed with OGT plus *Cp*OGA. b) TAB1^S395C^ and TAB1^S395C^ co-expressed with OGT plus *Cp*OGA crystals were fished, extensively washed in mother liquor and analysed for the presence of S-GlcNAc by Western blot. c) DSF profile of TAB1^S395C^ and TAB1^S395C^ co-expressed with OGT plus or minus *Cp*OGA. Each point in the graph represents the average of 6 replicates. Standard deviations are represented.

**Supplementary Figure S2.**
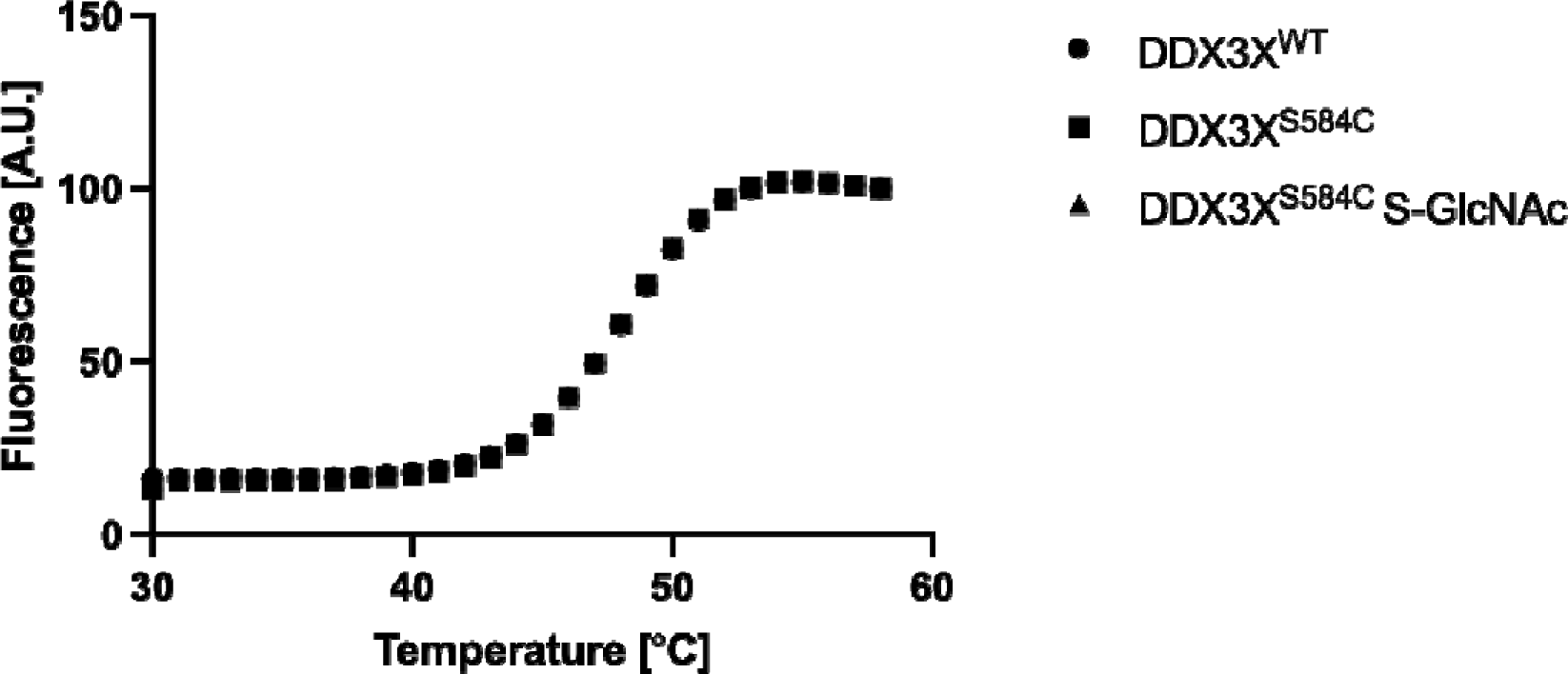
The thermal stability of DDX3X^WT^ and both un-modified and S-GlcNAcylated DDX3X^S584C^. The thermal stability of 1 mg/ mL of DDX3X was analysed by differential scanning fluorimetry, as described in the Materials and Methods. Fluorescence signals are shown relative to the maximal fluorescence signal detected for each replicate (n = 6). Due to the small error between technical replicates, error bars (standard deviation) are not visible.

**Supplementary Figure S3.**
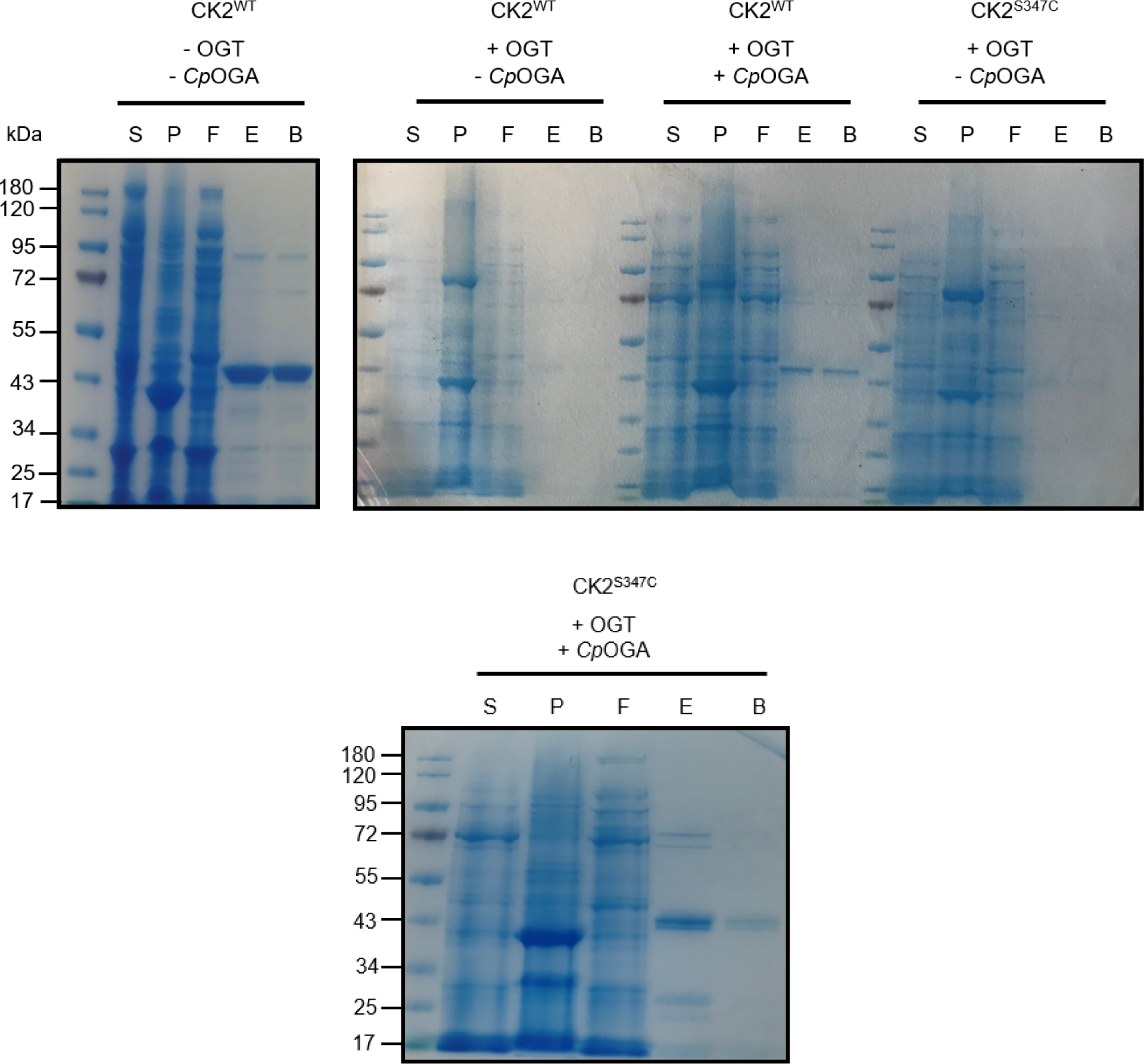
Comparison of soluble CK2α levels following co-expression with OGT with and without *Cp*OGA. Soluble (S) and pellet (P) fractions of IPTG-induced *E. coli* lysate, along with aliquots of flow-through (F), imidazole elutions (E), and Ni^2+^ beads post-elution (B), were analysed by SDS-PAGE and Coomassie staining to compare relative yields of CK2α in the presence or absence of OGT and *Cp*OGA.

**Supplementary Figure S4.**
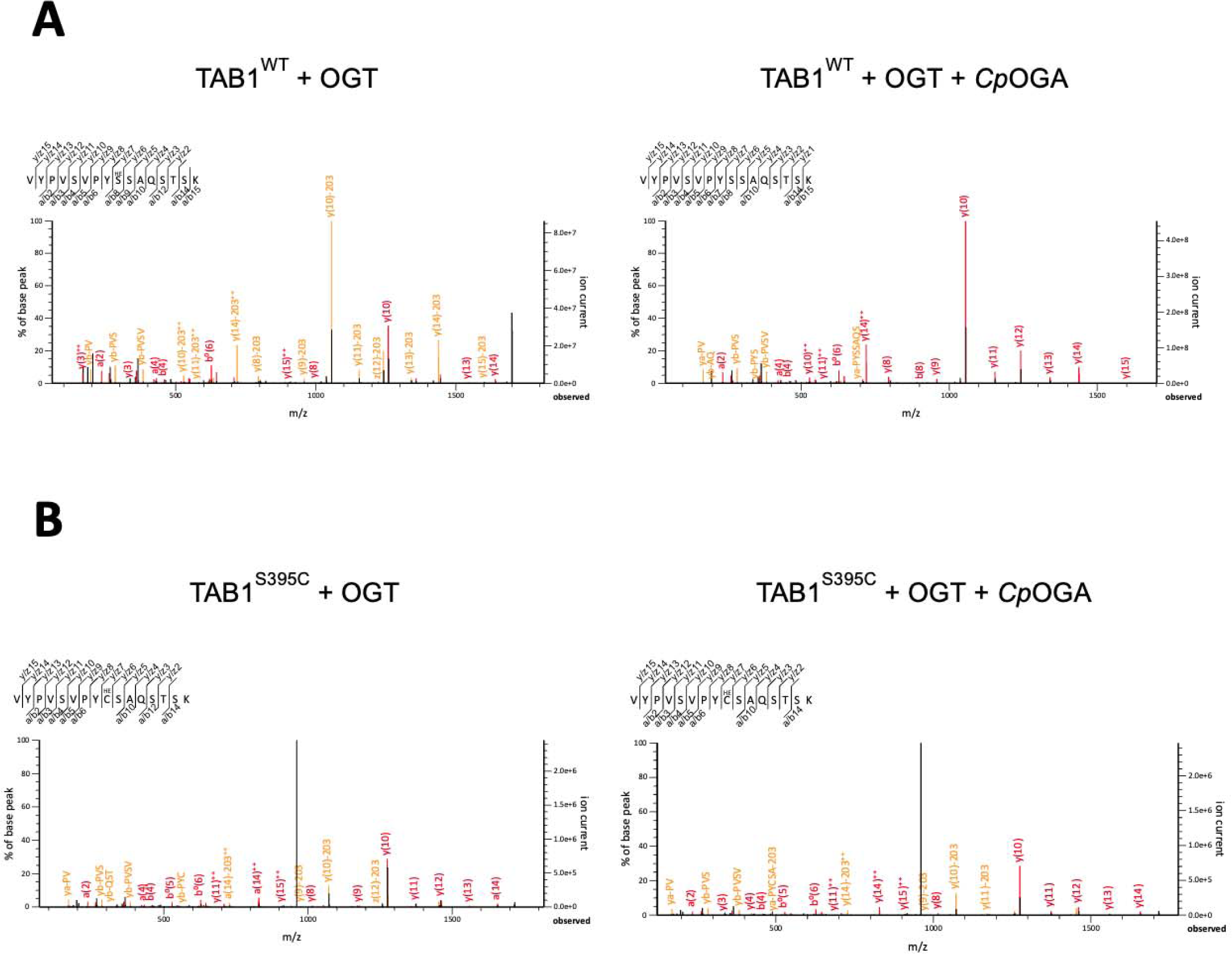
LC-MS/MS based identification of O- and S-GlcNAcylated TAB1 at position 395. a) Left hand panel: spectrum for the TAB1 VYPVSVPYSSAQSTSK peptide after co-expression with OGT only. Unambiguous identification of the O-GlcNAcylated residue was not possible using the Mascot search engine, which assigned equal probabilities to S4, S9, S10, S13, and S15. However, manual inspection of spectra reveals the presence of both y7 and y8 ion indicating position 9 (Ser395 in TAB1) is the likely modification site. Right hand panel: spectrum for the TAB1 VYPVSVPYSSAQSTSK peptide after co-expression of TAB1 with OGT/*Cp*OGA. Only the unmodified peptide was observed (see supplementary file 1). b) Left hand panel: spectrum for the TAB1^Ser395Cys^ VYPVSVPYCSAQSTSK peptide after co-expression with OGT only. As for a), S4, C9, S10, S13, and S15 were assigned equal probability of modification by Mascot. The absence of Cys alkylation and the presence of both the y7 and y8 ions indicate S-GlcNAcylation of at position 9 (Cys395 in TAB1^Ser395Cys^). Right hand panel: spectrum for the TAB1^Ser395Cys^ VYPVSVPYCSAQSTSK peptide after co-expression of TAB1^Ser395Cys^ with OGT and *Cp*OGA. As for the left hand panel of b), the absence of Cys alkylation and the presence of y7 and y8 ions indicates mono-S-GlcNAcylation at position 9 (Cys395 in TAB1^Ser395Cys^).

**Supplementary Table 1.**
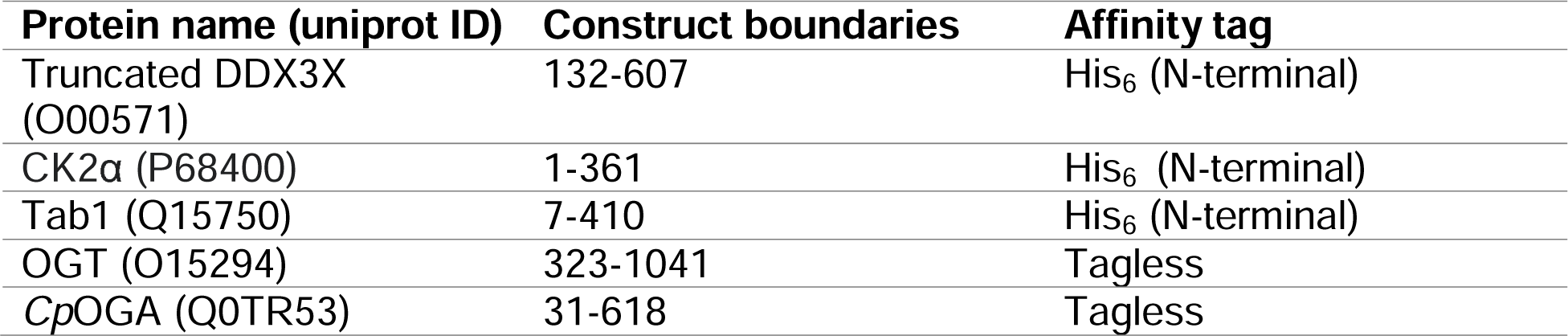
Protein constructs used in this study. Uniprot IDs of proteins are provided, along with construct boundaries. The affinity tag used for each construct is also provided. All tagged constructs are linked to their respective affinity tags via a PreScission protease cleavage site

| **Protein name (uniprot ID)** | **Construct boundaries** | **Affinity tag** |
| --- | --- | --- |
| Truncated DDX3X (O00571) | 132-607 | His<sub>6</sub> (N-terminal) |
| CK2α (P68400) | 1-361 | His<sub>6</sub> (N-terminal) |
| Tab1 (Q15750) | 7-410 | His<sub>6</sub> (N-terminal) |
| OGT (O15294) | 323-1041 | Tagless |
| CpOGA (Q0TR53) | 31-618 | Tagless |

